# High-throughput isolation and sorting of gut microbes reduce biases of traditional cultivation strategies

**DOI:** 10.1101/759969

**Authors:** William J. Watterson, Melikhan Tanyeri, Andrea R. Watson, Candace M. Cham, Yue Shan, Eugene B. Chang, A. Murat Eren, Savaş Tay

## Abstract

Traditional cultivation approaches in microbiology are labor-intensive, low-throughput, and often yield biased sampling of taxa due to ecological and evolutionary factors. New strategies are needed to enable ample representation of rare taxa and slow-growers that are outcompeted by fast-growing organisms. We developed a microfluidic platform that anaerobically isolates and cultivates microbial cells in millions of picoliter droplets and automatically sorts droplets based on colony density. We applied our strategy to mouse and human gut microbiomes and used 16S ribosomal RNA gene amplicons to characterize taxonomic composition of cells grown using different media. We found up to 4-fold increase in richness and larger representation of rare taxa among cells grown in droplets compared to conventional culture plates. Automated sorting of droplets for slow-growing colonies further enhanced the relative abundance of rare populations. Our method improves the cultivation and analysis of diverse microbiomes to gain deeper insights into microbial functioning and lifestyles.

## Introduction

Culture-independent surveys of naturally occurring microbial populations through marker gene amplicons and shotgun metagenomes have revealed intriguing associations between the gut microbial communities and human health [1, 2]. However, inferring the taxonomic composition or functional potential of complex gut microbiomes does not reveal mechanistic underpinnings of observed associations [3-5]. Microbial cultivation is an essential first step to address some of these shortcomings as this strategy enables the recovery of complete reference genomes [6], accurate identification of taxonomy and functional potential of new strains [7, 8], and validation of causality by perturbation experiments [5]. Microbial cultivation is currently experiencing a pronounced revival [7-11], yet the majority of cultivation efforts require arduous manual picking of thousands of colonies, impeding the efforts to harmonize discoveries that emerge from ‘omics strategies with downstream mechanistic investigations in the rapidly advancing field of microbiome research.

In recent years, numerous cultivation strategies have been developed to increase the throughput in isolating and studying gut-associated bacteria. For example, a recent well plate-based growth experiment screened 96 phylogenetically diverse human gut-associated bacterial strains across 19 media and determined their nutritional preferences and biosynthetic capabilities [12]. Another study that relied on ‘SlipChip’ [13], a microfluidic device that can isolate hundreds of microbial cells and enable targeted cultivation, successfully recovered an organism that was a member of the genus *Oscillibacter* [14]. Until this study, *Oscillibacter* had been one of the “most wanted taxa” from the human gut [15], a list of uncultivated yet highly prevalent taxa from the Human Microbiome Project [16]. Biomimetic devices represent another active area of research [17]. For instance, the “gut-on-a-chip” offers a controlled microfluidics platform which mimics the physical and functional features of the intestinal environment and enables complex *in vitro* chemical gradients and multicellular interactions [18, 19] that can establish stable co-culturing of complex bacterial populations [20]. Although these techniques increase the throughput in isolating and manipulating gut organisms as compared to agar culture, their throughput is insufficient for isolating rare organisms among the thousands of gut-associated species or performing large-scale perturbation experiments.

Droplet microfluidics offers a promising alternative for high-throughput anaerobic cultivation. The aqueous droplets, with typical volumes ranging from picoliters to nanoliters, are generated and manipulated with an oil phase in microfluidic channels. An extensive arsenal of droplet microfluidic tools has been developed for use in standard aerobic lab environments, where oxygen is present [21]. For instance, droplets can be generated and sorted at rates exceeding 10 kHz [22], reagents can be added by pico-injection or droplet merging [23, 24], and the droplets can further be stored in a regular array and retrieved for downstream applications [25, 26]. Droplets also eliminate a major bottleneck of conventional broth and agar culture: the overgrowth of fast-growing populations over slow-growers. In particular, the stochastic isolation of individual bacterial cells in discrete droplets prior to cultivation eliminates the competition that favors fast-growers and yields more accurate representation of the distribution of microbial cells from the input sample. For instance, Jiang et al. (2016) isolated environmental soil-associated bacteria in droplets and found an increase in the diversity of taxa with an increased representation of rare organisms [25]. Villa et al. (2019) recently cultivated human gut microbes in thousands of nanoliter droplets to characterize metabolic variation in polysaccharide-degrading gut bacteria and analyzed their growth kinetics [11]. Finally, bacteria confined within droplets reach a critical threshold concentration of quorum sensing molecules faster than would occur in bulk culture (due to the inherently small droplet volume), which can lead to improved growth in certain culture media [27]. These three advantages afforded by droplet microfluidics would thus present an ideal technology for improving the speed and efficiency of traditional anaerobic culture provided droplet manipulation techniques could be extended to anaerobic environments.

Here, we present an end-to-end platform for high-throughput automated isolation, cultivation, and sorting of anaerobic bacteria in microfluidic droplets. The technology is comprised of droplet microfluidic devices operated inside of an anaerobic chamber and an automated rapid image processing system. We first demonstrated our technology’s ability by isolating and cultivating an 8-species model mouse gut community, the Altered Schaedler’s Flora. We observed that all 8 species grew in the droplets, including the extremely oxygen sensitive species, and the droplets prevented the competitive overgrowth of a fast-growing species over slow-growing species. We then cultivated human fecal bacteria across 3 different rich media using our microfluidic platform and compared the growth to that on conventional agar plates. The droplet cultivation featured several advantages including an increase in community richness over agar plates by 37% up to 410%, depending on the medium, and an enhancement in the cultivation of low abundance (<1%) strains in raw stool. We also found a reduction in the variability of community composition across different media, and the droplets enabled cultivation of strains from the clinically important genus, *Bacteroides*, several of which did not grow on plates within the experiments conducted here. Further, sorting droplets based on colony density led to a further enhancement of low abundance strains in the sorted fraction. Finally, to ensure that our droplet technology is compatible with traditional microbiology workflows, we streaked sorted droplets onto an agar plate and observed standard colony growth. In total, our droplet-based anaerobic isolation and cultivation technology has the potential to increase the throughput and success rate of recovering pure strains from human gut-associated samples.

## Results

### Isolation, culture, and sorting of anaerobes in a high-throughput droplet microfluidic system

We developed an array of droplet microfluidic technologies for the high-throughput cultivation and manipulation of anaerobic microbial communities (Fig. 1). The microfluidic devices are housed within an anaerobic chamber along with a microscope, syringe pumps, a high frame rate camera, electrodes, and an incubator (Fig. 1a). A computer external to the anaerobic chamber controls the camera, syringe pumps, and electrodes (via a voltage amplifier also external to the chamber). The droplets are generated at a flow focusing junction from liquid culture medium into oil (Fig. 1b). The bacteria are stochastically encapsulated in the droplets (∼65 – 115 pL) by diluting fecal cell suspensions or liquid broth cultures in the medium so that ∼20% to 30% of droplets initially contain only one live bacterial cell according to Poisson statistics. The droplet emulsion is then placed within an incubator at 37 °C and the isolated viable strains can clonally replicate within a droplet provided the strain can grow within the environmental conditions (Fig. 1c). Since each colony is isolated within a confined droplet, the slow-growing species avoid the competitive overgrowth of fast-growing species – which often occurs in traditional broth or Petri agar cultivation.

**Figure 1.**
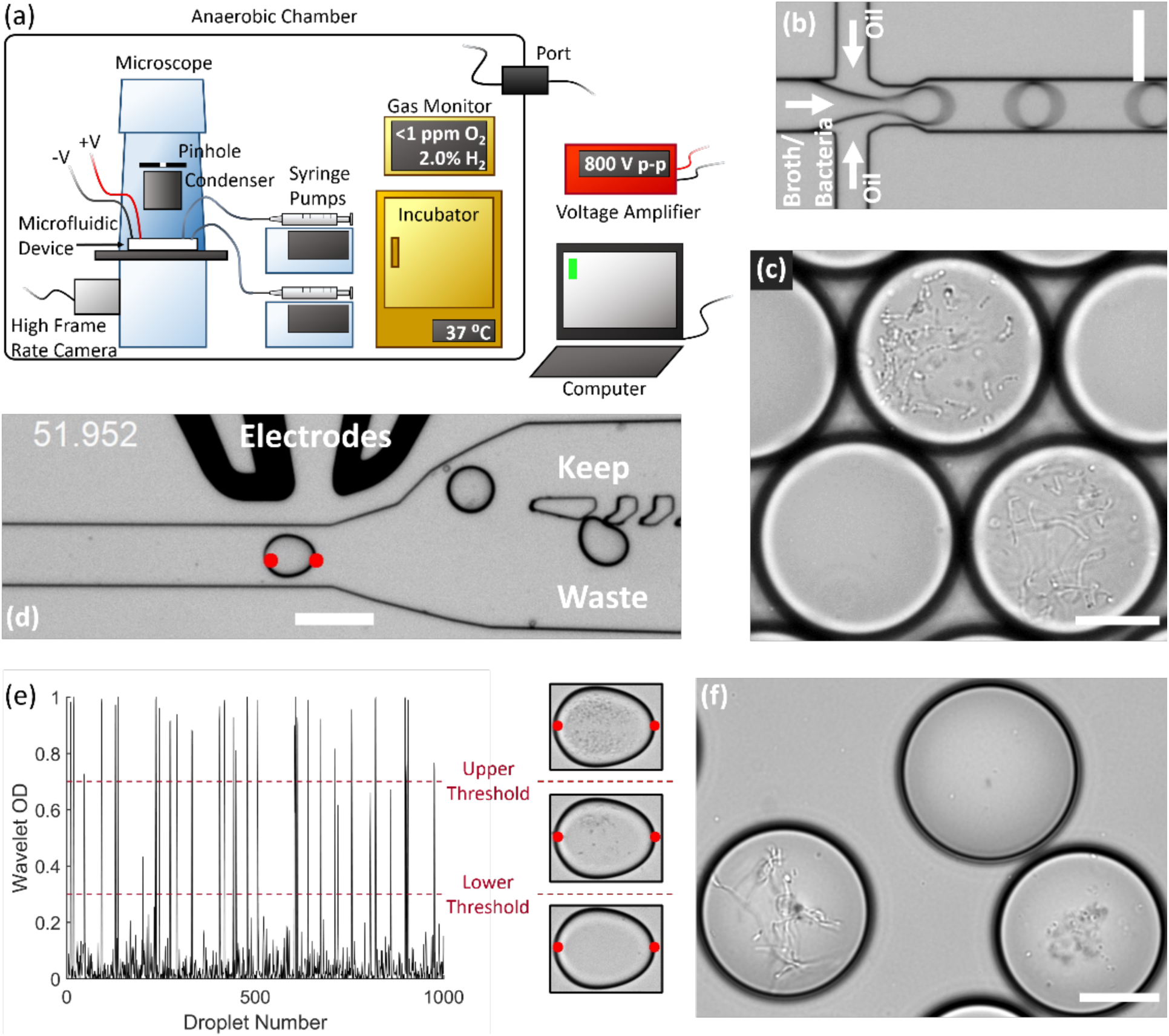
Experimental setup for isolation and culture of gut anaerobes in microfluidic droplets. (a) The experimental setup for isolating, culturing, and sorting anaerobic bacteria in microfluidic droplets consists of a microscope, microfluidic devices, a high frame rate camera, syringe pumps, an incubator, and electrodes all contained within an anaerobic chamber. The computer controls the syringe pumps, high frame rate camera, and electrodes (via a voltage amplifier). The equipment power and control wires are introduced to the anaerobic chamber through sealed rubber ports to strictly maintain the anaerobic conditions within the chamber. (b) Single bacteria cells are isolated in droplets containing anaerobic culture medium and the resulting emulsion is cultured inside the incubator. (c) An example of human gut bacteria isolated and cultivated inside droplets. (d) Droplets are sorted by optical detection and subsequent deflection via dielectrophoresis near a sorting junction. Specifically, droplets with bacterial colonies which meet a certain thresholding criteria were determined using image analysis (region between the red dots), and these droplets were deflected into the “keep” path by actuating an on-chip electrode while sending the remaining droplets to waste. (e) The colony density measured by image analysis (Wavelet OD) for 1000 successive droplets. Droplets with a dense colony, a sparse colony, and empty droplets (no colony) are represented by a wavelet OD value above an upper threshold, between an upper and lower threshold, or below a lower threshold, respectively. (f) Two slow-growing human gut-associated bacteria colonies (bottom left and bottom right) grown in droplets after sorting and a false positive empty droplet (top). Scale bar in (b) and (d) is 100 μm and scale bar in (c) and (f) is 20 μm.

We also developed an image-based sorting algorithm and microfluidic control system for sorting bacterial colonies in droplets based on the colony density (Fig. 1d). Importantly, image-based droplet sorting does not require fluorescent strains or reporters and therefore has broad applicability in processing environmental samples [28]. The high-frame rate camera along with a custom LabView code automatically detects the droplet approaching the sorting junction and performs a wavelet-based image analysis of an optical density-like measurement, which we termed the Wavelet OD (see Methods). If the Wavelet OD satisfies an empirically-defined thresholding criteria, the computer will actuate the electrodes via a voltage amplifier and deflect the droplet bacteria colony into the “keep” path (Fig. 1d). The Wavelet OD value varies between 0 (empty droplets) and 1 (droplets with a very dense colony). Droplets were sorted at a rate of ∼30 Hz. Slow-growing colonies were retrieved by sorting droplets with a Wavelet OD within a lower and upper threshold value (Fig. 1 e-f).

### Validation of an 8-species model mouse gut community, the Altered Schaedler’s Flora

The Altered Schaedler’s Flora, ASF, is an 8-species model bacterial community representative of the normal mouse gut microbiota [29]. ASF is an important and widely studied gnotobiotic mouse model used for understanding microbiota-host dynamics in both health and disease. The ASF community contains 2 facultative anaerobes, *Lactobacillus intestinalis* (ASF 360) and *Lactobacillus murinus* (ASF 361), 2 anaerobes, *Mucispirillum schaedleri* (ASF 457) and *Parabacteroides goldsteinii* (ASF 519), and 4 extremely oxygen sensitive anaerobes, *Clostridium sp*. (ASF 356), *Eubacterium plexicaudatum* (ASF 492), *Pseudoflavonifactor sp*. (ASF 500), and *Clostridium sp*. (ASF 502) [29, 30].

We investigated whether pure ASF strains as well as a mixed community of ASF strains from a fecal pellet could be cultured anaerobically in droplets. Pure ASF strains were cultured from stock solutions in liquid BHIS-ASF broth, then diluted and encapsulated in droplets using our microfluidic platform. We observed that all 8 ASF strains were successfully cultured in the droplets (Fig. 2a and Supp. Fig. 1). We obtained ASF fecal pellets (Taconic Biosciences) and created stock solutions of the mixed community cell suspension. The ASF feces contained 7 of the 8 strains (Supp. Fig. 2); ASF 360 was not detected. We performed 3 biologically independent cultures of the ASF community in broth and droplets (Fig. 2b-c). When we cultured the ASF community in liquid BHIS broth, ASF 361 outgrew the other strains. We note that ASF 361 is parasitic towards ASF 519 [31], which may explain the dominance of ASF 361 in broth culture. On the other hand, culturing within droplets confined the growth of ASF 361 and allowed a larger relative abundance of ASF 519 (34%) and ASF 457 (0.96%) to grow (Fig. 2c). Contrary to the pure ASF strain cultures, the four extremely oxygen sensitive species from the ASF fecal pellets exhibited extremely limited to no growth in either bulk liquid culture or the droplets, likely due to oxygen exposure during collection and shipping. Finally, we observed similar colony morphologies of ASF 361, 457, and 519 between the pure strain and mixed community droplet cultures, with the exception of spore forming communities from the mixed community sample (Fig. 2b).

**Figure 2.**
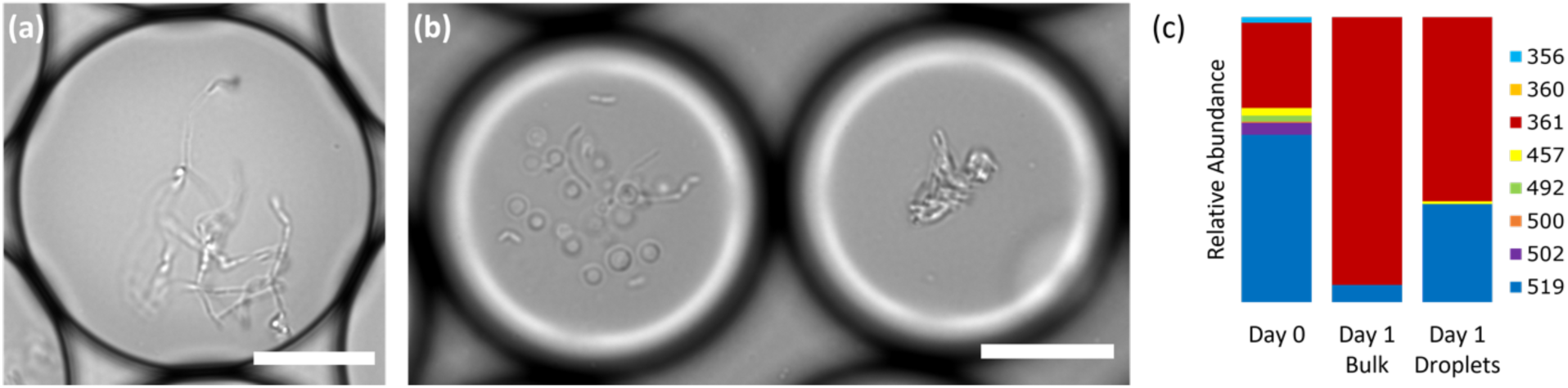
Droplet-based culture of the ASF, a model mouse gut community. (a) The extremely oxygen sensitive anaerobe, ASF 502, grown in microfluidic droplets after 20 h from a pure species culture. (b) Single ASF species were isolated in droplets using a mixed community of bacterial cell suspension obtained from an ASF fecal pellet and cultivated for 24 h. The colony morphology in the right droplet is visually similar to ASF 361 grown from pure strain cultures, while the left droplet has a spore-forming community which was not observed in the pure strain cultures. (c) Relative abundance of ASF strains in the fecal cell suspension before cultivation (Day 0), grown in bulk culture from the fecal suspension (liquid broth) for 1 day, and grown in droplets from the fecal suspension for 1 day. The relative abundance for bulk and droplet cultures are averaged over 3 independent experiments. Scale bars in (a) and (b) are 20 um.

### Cultivation of human stool microbiota in droplets enhances richness and abundance of rare taxa

In order to explore the growth potential of human gut bacteria in microfluidic droplets, we cultivated bacteria in rich medium droplets (Fig. 1c and Supp. Video 1) and compared the growth to that on conventional agar. In particular, we collected a single human fecal sample, created glycerol stock cell suspensions, and cultured the bacteria in droplets for up to 2 days or on agar plates for up to 7 days. The 3 rich media utilized were Brain Heart Infusion Supplemented (BHIS), Gut Microbiota Medium (GMM), and Yeast Casitones Fatty Acid (YCFA). After cultivation, we extracted the genomic DNA from plate scrapings or by breaking the pooled droplet emulsion. The samples were sequenced using paired-end 16S rRNA sequencing on the Illumina platform with primers targeting the V4 region. To infer highly resolved microbial community structures in our amplicon data, we used Minimum Entropy Decomposition (MED) [32], which uses Shannon entropy to identify highly variable nucleotide positions among amplicon sequences and iteratively decomposes a given sequencing dataset into oligotypes, or “amplicon sequence variants” (ASVs), in which the entropy is minimal. The single-nucleotide resolution afforded by this strategy allows the identification of closely related but distinct taxa, better explaining micro-diversity patterns that may remain hidden otherwise [33, 34]. Our data revealed no significant variation in the community composition between biological replicates or cultivation time for a given culture method (droplets or agar plates) and media (Supp. Fig. 3). Irrespective of the media and cultivation time, the community richness (number of detected ASVs) in droplets was larger than that on agar plates (Fig. 3a-b, *p* < 0.005, Mann-Whitney *U* test). In particular, droplets enabled an increase in richness between 37% (BHIS) and 410% (YCFA). The community diversity, measured by the Shannon index, increased in BHIS and YCFA droplets over their corresponding agar plates, but not in GMM (Fig. 3c). The most abundant ASV in all droplet samples, except 2-day GMM droplets, belonged to the genus *Hafnia* and had a mean abundance of 23% averaged across media and cultivation time (Supp. Fig. 4). The droplets also featured a similar taxonomic composition at the family level across the three media, whereas the agar plate cultures more drastically differed from each other and from the input sample (Fig. 3d and Supp. Fig. 5).

**Figure 3.**
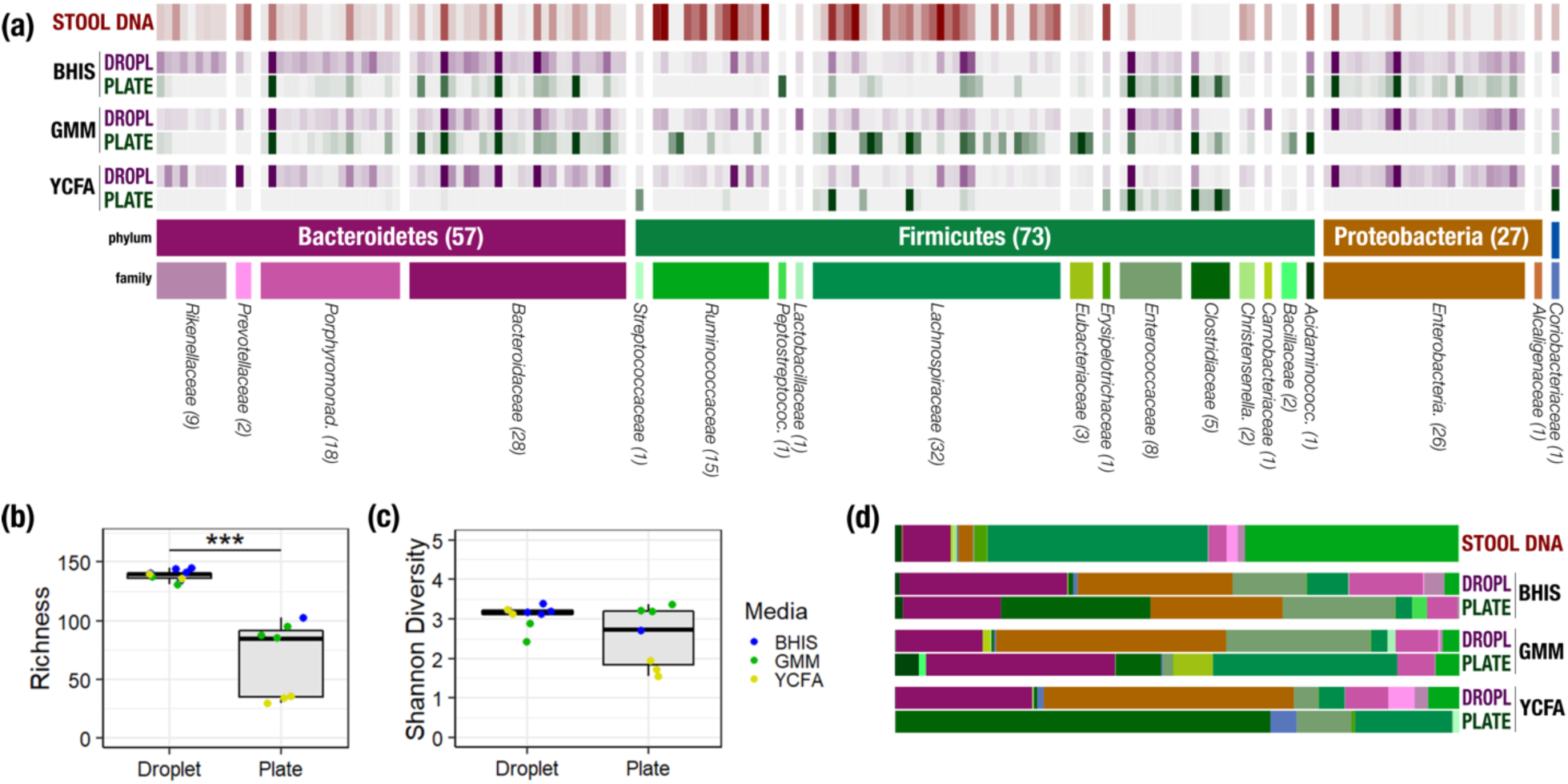
Comparison of human stool bacteria cultured on agar plates versus in droplets. (a) The relative abundance for each ASV organized by phylum and family is plotted for the raw stool and for representative droplet and plate cultures in each media. The dark bars indicate a higher relative abundance. (b) The community richness averaged over cultivation time and media is increased in droplets over plates (*p* < 0.005) but (c) the Shannon diversity is not. (d) Family-level relative abundance for representative droplet and plate cultures. The community composition between droplet cultures of different media is more similar than between plates.

**Figure 4.**
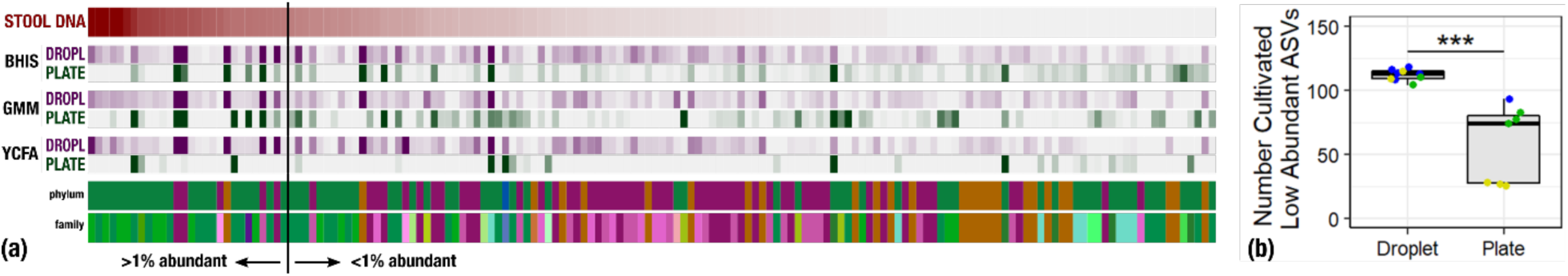
Droplets improve the cultivation of low abundant organisms. (a) The relative abundance per ASV organized by relative abundance in raw stool. The phylum and family colors correspond to the labels shown in Fig. 3a. (b) The number of cultivated low abundant ASVs in the raw stool sample (<1%, total of 130 ASVs) averaged over cultivation time and three different media is increased in droplets over plates.

**Figure 5.**
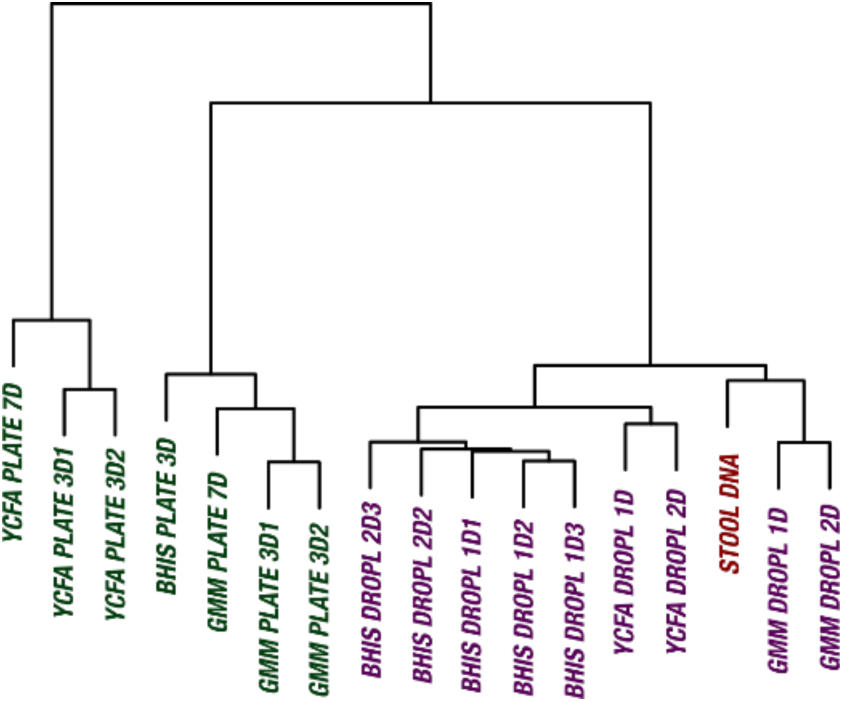
The culture of stool in droplets enables cultivation of clinically relevant *Bacteroides spp*. which did not grow on plate cultures. Hierarchical clustering of *Bacteroides* oligotypes across 7 plate cultures and 9 droplet cultures reveals a stronger association of droplet cultures towards the raw stool as compared to plates.

One of the general bottlenecks of plate-based cultivation efforts is to isolate organisms that are rare in the input sample, because abundant taxa are often over-represented in plates [8]. Our data showed that droplets were able to grow a larger number of organisms that were low-abundance (<1%) in the original stool sample based on 16S rRNA gene amplicons (Fig. 4, *p* < 0.005, Mann-Whitney *U* test). In particular, out of 130 ASVs that were <1% abundant in the original stool sample, 112 ASVs from droplets and 58 ASVs from plates, averaged over cultivation time and media, were detected. Next, we investigated how the composition of closely related taxa that resolve to the same taxonomic group in our cultivation efforts compared to their composition in the stool. For this, we performed an oligotyping analysis on all sequencing reads that matched to a single taxon, *Bacteroides*, the most abundant genus in our dataset and one of the most variable genera across individuals [35]. Hierarchical clustering of our samples based on the distribution of *Bacteroides* oligotypes revealed that the composition of *Bacteroides* populations measured by the 16S rRNA gene amplicons in droplet-based cultures were more similar to those in raw stool than plate-based cultures, as they clustered closer to the stool sample (Fig. 5, *p* < 0.005, multiscale bootstrap resampling). This indicates the influence of growth biases associated with plate-based cultivation was lessened in droplet-based cultures regardless of the medium, and a larger fraction of *Bacteroides* populations were accessible through droplets (Fig. 5).

### Sorting slow growing organisms in droplets further amplified the abundance of rare taxa

Relatively slow growth rate is one possible explanation for the apparent low abundance of any given taxon within a sample. To investigate whether we could increase the relative abundance of ASVs which were <1% abundant in the raw stool sample, we automatically sorted droplets based on the colony density (Supp. Video 2). We performed two independent sorting experiments using human stool samples grown in BHIS droplets to keep only low-density colonies. The false positive sorting rate was low, with at most 8% of droplets incorrectly sent into the keep path. Sorted droplet cultures resulted in a shift in community composition (*p* < 0.005, Kolmogorov-Smirnov test), with a noticeable change in the abundance of the top 20 ranked ASVs (Fig. 6a). Next, we investigated which ASVs were amplified from <1% in raw stool to >1% in unsorted and sorted droplets. The average number of ASVs amplified from the <1% to >1% condition in unsorted BHIS droplets was 6.5 out of the 158 ASVs detected in total, while sorting increased the average number of amplified ASVs to 12.5 out of 158 (Fig. 6b), thereby indicating that droplet sorting based on optical density can provide some preference in amplifying low abundance taxa. Additionally, we note that sorted samples amplified taxa across a wider range of the phylogenetic tree than unsorted samples. Finally, to ensure that bacteria remain viable after droplet cultivation, we streaked sorted droplets from one experiment onto an agar plate and cultured the bacteria on the plate for 2 days. We observed colony growth and random picking of 24 colonies recovered species from the genera *Hafnia* (12/24), *Enterococcus* (8/24), and *Bacteroides* (4/24) (Supp. Fig. 6).

**Figure 6.**
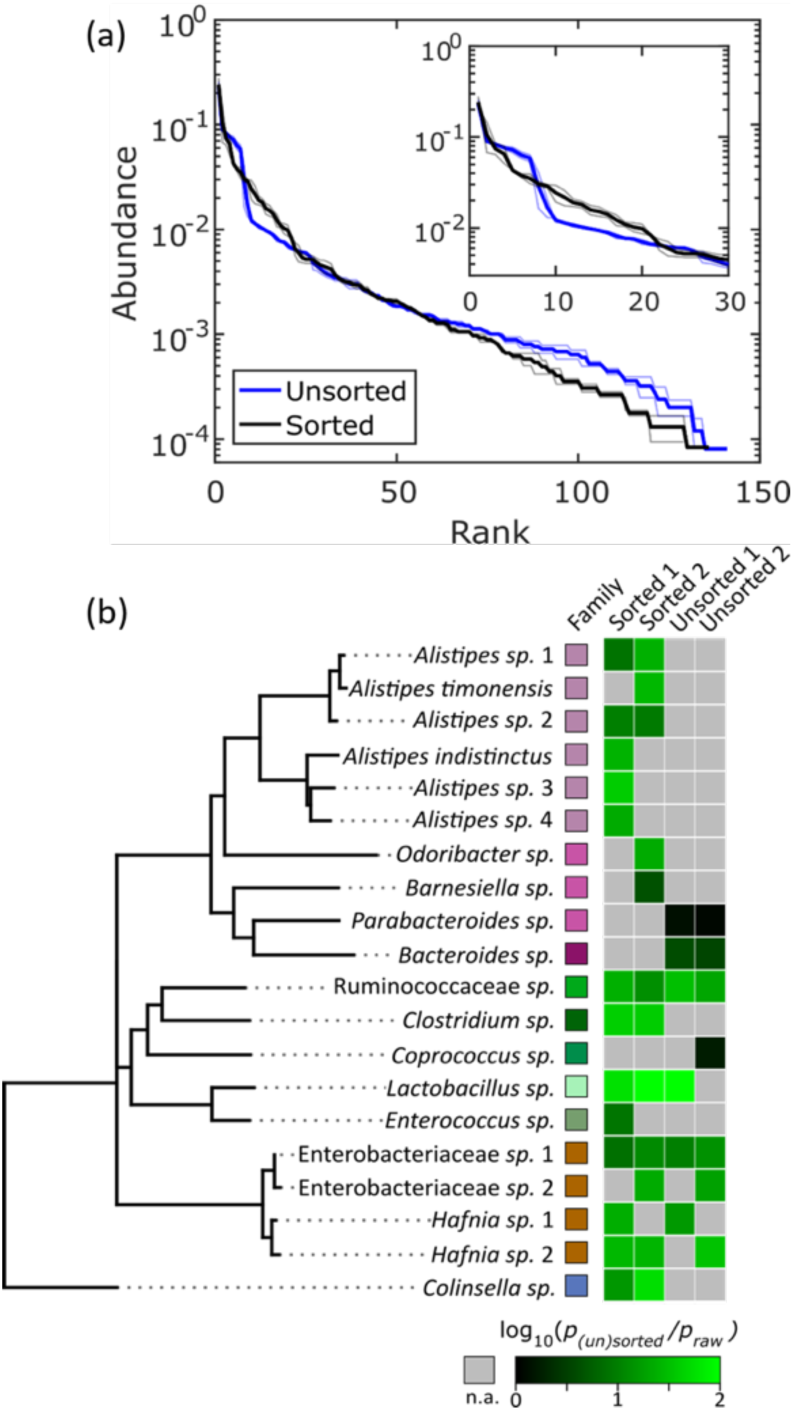
Isolation of slow-growing species from human stool microbiota. (a) The rank-abundance curves show that sorting based on colony density changes the overall community composition. A zoomed in portion of the rank-abundance curve is shown in the inset. (b) Phylogenetic tree of ASVs which were <1% abundant in raw stool but were increased to >1% in at least one sorting experiment. The ratio of amplification for strains amplified above the 1% limit is depicted in the heat map. N.A. (shown in gray) indicates ASVs which were not amplified from <1% to >1%. The legend for the family color labels is depicted in Fig. 3a.

## Discussion

Here, we developed a droplet-based microfluidic platform for isolating, cultivating, and sorting mouse and human gut-associated anaerobic bacteria. The technology consists of droplet generating and droplet sorting microfluidic devices operated inside an anaerobic chamber along with automated sorting based on colony density. We first prototyped our method using a mouse model gut community, the Altered Schaedler’s Flora, and subsequently explored the cultivation of bacteria from a human stool sample.

The droplet-based culture provides several key benefits over traditional agar cultivation. First, droplets offer a high-throughput platform for culturing, manipulating, and monitoring bacterial colonies. In aerobic systems, droplet microfluidics has enabled significant progress in fields ranging from pathogen detection, to antibiotic susceptibility testing, to strain engineering [21]. Here, we demonstrated that droplet generation, cultivation, and sorting can be extended to anaerobic systems, potentially paving the way towards improved throughput in studying gut microbiota. Second, parasitism, amensalism, and competition are eliminated between strains since each colony is isolated in its own droplet. For instance, in broth co-culture ASF 361 is parasitic towards ASF 519, leading to an increased abundance of ASF 361 and a decreased abundance of ASF 519 [31]. Here, we observed ASF 361 dominated the broth culture of the mixed ASF community. However, the droplets confined the growth of ASF 361 and allowed ASF 457 and ASF 519 to reach a higher relative abundance. Finally, the concentration of quorum sensing molecules can increase in droplets faster than bulk culture due to the inherently small volume of each droplet [27]. Potentially, quorum sensing mediated growth could explain the predominant abundance of *Hafnia spp*. in droplets across all three rich media, since the growth dynamics of at least some strains of *Hafnia alvei* are modulated by quorum sensing [36].

These benefits of the anaerobic droplet technology translated into significant improvements in the cultivated community representation of the gut bacteria. In particular, we found: (1) the community richness was increased by up to 410% in droplets over agar plates, (2) the droplet-based culture reduced the variation between media in the cultivated taxonomic composition, (3) the number of low abundance strains from raw stool was also increased in droplets, (4) the droplet-based culture enabled cultivation of clinically relevant *Bacteroides* which were not detectable on plate cultivations in our study, and (5) sorting based on colony density further amplified the abundance of certain low abundance strains thereby facilitating the isolation of rare species. Moreover, the sorted bacteria remained viable and multiple taxa grew on agar plates following streaking of the droplets. Given that the ultimate goal is to isolate pure strains, our technology can therefore help facilitate isolation by boosting the community richness and initializing the growth of certain strains across a wide range of taxa before applying more conventional microbiological methods.

We envision that several further refinements to our droplet-based cultivation strategy could address limitations of the current study. First, in order for anaerobic droplet cultivation to be widely utilized among microbiologists, the droplet generating and sorting technologies must be easily integrated into standard biology workflows. The methods presented here are likely immediately transferrable for isolation and cultivation of anaerobic bacteria in droplets, since microfluidic droplet generating devices are now commercially available [37]. However, anaerobic droplet sorting is technologically more involved and will likely require commercial development of an anaerobic droplet sorter before it is widely adopted. Second, in this study, we isolated single living bacterial cells into droplets in order to prevent interspecies competition. However, isolation inhibits the growth of organisms that rely on other microbial or host cells, such as Saccharibacteria (formerly TM7) [38], or obligate endosymbionts, such as *Wolbachia* [39]. In droplets, co-encapsulation of two cross-feeding auxotrophic strains into a single droplet can induce growth, whereas growth will not occur when only one auxotroph is present within a droplet [40, 41]. Future studies could stochastically co-encapsulate multiple gut bacteria into droplets (by increasing the loading cell density during droplet generation) and investigate the resulting growth dynamics. Finally, here we loaded droplets with three different rich media in order to broadly enrich the cultivated community representation across taxa. However, some bacteria prefer defined media over rich media [12] while others utilize surface features such as hydrophobicity, roughness, and surface chemistry to form biofilms and proliferate [42]. Further enrichment of anaerobic organisms within our droplet platform could be achieved by incorporating droplet generation with defined media [11], combinatorially generated gradients of media [43], or varying the droplet surfactant chemistries to improve biofilm formation [44].

Despite large-scale cultivation efforts employing the manual picking of tens of thousands of colonies [7-10], a significant fraction of gut microbiota remains uncultured. In fact, recent large-scale analyses that reconstructed population genomes directly from gut metagenomes indicate that our current culture collections lack 40% to 50% of genera encompassing ∼2,000 species [45-47]. Many of these organisms may be rare, present in only a small proportion of individuals, and/or not readily amenable to culture on standard agar plates, further compounding the difficulty in obtaining isolates using conventional microbiological methods.

High-throughput anaerobic droplet cultivation has a significant potential to accelerate the study of gut bacteria and the isolation of presently uncultivated species. For instance, in parallel to our work here, Villa et al. (2019) recently cultivated human stool microbes in droplets and screened carbohydrate utilization of bacteria from nine stool donors in order to determine the inter-individual variation in metabolic utilization [11]. Here, we demonstrated that our technology enabled an enrichment of low abundance organisms and reduced the variation of the cultivated taxonomic composition between media. Our technology could therefore facilitate an increased recovery of strains from human samples, highlighting the potential for anaerobic droplet microfluidics in elucidating the function of complex microbiomes such as in the gut human health and disease.

Our approach for isolation, cultivation, and sorting of gut microbiota in droplets enriched the representation of taxa across bacterial phyla, including organisms which are rare and/or slow-growing. In particular, while recovering bacteria using traditional cultivation techniques is often biased towards abundant or fast-growing organisms, isolation of bacteria in droplets inhibited interspecies competition resulting in an enriched abundance of rare and slow-growing organisms. Bacteria remained viable throughout the droplet cultivation and sorting processes suggesting that our anaerobic droplet technology is compatible with traditional downstream microbiology techniques. Going forward, our technology could facilitate overcoming difficulties in traditional plate-based cultivation and pave the way for rapid recovery of novel strains in complex systems such as the human gut microbiome.

## Methods

### Microfluidic device fabrication

The microfluidic droplet generation and droplet sorting devices were fabricated using soft lithography techniques. The device architectures were adapted from Mazutis et al. (2013) [48]. Briefly, we first fabricated molds from the negative photoresist, SU-8 3050, on 4” silicon wafers. The height of both the droplet generation and sorting devices was 50 μm. We then poured a 10:1 ratio of PDMS (RTV 615) parts A to B onto the mold, degassed the PDMS, and cured at 80 °C for at least 1 hour. Next, we removed the cured PDMS from the mold, punched holes for the inlets, outlets, and electrodes, and plasma bonded the PDMS to either a glass slide (for droplet generating devices) or a glass slide with a conductive indium-tin oxide on the rear side (for droplet sorting devices, Delta Technologies - Part No. CG-811N-S207). To increase the microchannel hydrophobicity, we coated the microchannels with Aquapel (Pittsburgh Glass Works) followed by Fluorinert FC-40 (Sigma). Finally, for droplet sorting devices, we created the electrodes by placing the microfluidic device on a 90 °C hotplate, flowing a low melting temperature solder (Indium Corporation of America, 51% In/32.5% Bi/16.5% Sn) into the electrode holes, and connecting the solder to standard wires.

### Microfluidic device control system

The syringe pumps, high-frame rate camera, and electrodes are controlled through custom written LabView code. We generated droplets using two syringe pumps (Harvard Apparatus Pump 11 Pico Plus Elite) which controlled the liquid and oil (Bio-Rad Droplet Generation Oil for EvaGreen) flow rates. We used two separate droplet generating devices here (see supplementary CAD files) with flow rates specified in the Supplementary Table, Experiment Info. The droplet volumes ranged from ∼65 – 115 pL. In a typical experiment, approximately 0.5 – 1 mL of droplets were generated in approximately 20 min – 1 hr, depending on the droplet generating device. For droplet sorting, the droplet reinjection flow rate was set to 20 μL/hr and the oil phase for droplet spacing was set to 180 μL/hr. The microfluidic devices were monitored using an inverted microscope (Nikon Ts2R) under 4x and 10x magnification. A high-frame rate camera (Basler acA640-750um) captured 672 × 360 pixel images at a rate of 925 Hz and the exposure time was set to 59 μs per frame. Our LabView code automatically analyzed each droplet near the sorting junction and made a sorting decision based off the wavelet OD (see droplet image analysis). Droplets which satisfied the sorting conditions were sent into the keep path by actuating the electrodes (Fig. 1d). The remaining droplets flowed down the waste path. The sorting rate was ∼30 Hz. The electrodes were actuated by outputting a true decision to an NI-DAQ 6211 which set the analog out to the desired voltage. The analog output voltage is then amplified (TREK Model 2220) at 200 V/V. The electrode actuations used a 10 kHz, 800 V p-p, 30% duty cycle square wave which was activated for 10 ms. An upper limit on the false positive sorting error, *ε*, is given by *ε* ≤ [*f* ^*un*^ (1-*f* ^*s*^)] / [*f* ^*s*^ (1-*f* ^*un*^)], where *f* ^*un*^ is the fraction of slow-growing colonies in the unsorted sample and *f* ^*s*^ is the fraction of slow-growing colonies in the sorted sample (i.e., the droplets sent into the keep path). The estimate on *ε* is an equality when the false negative rate is zero. However, we did not count the slow-growing fraction in the waste stream and therefore the actual false positive rate is likely lower than the upper limit.

### Droplet image analysis

We sorted bacterial colonies in droplets based on an optical density-like measurement, which we termed the Wavelet OD. Custom LabView code first located each droplet as it approached the sorting junction by detecting the droplet edges along the center of the channel (i.e., the red dots in Fig. 1d). The interior region of each droplet (∼60 × 80 pixels) was then analyzed using a discrete wavelet frame decomposition from the LabView function IMAQ Extract Texture Feature VI [49]. We optimized the wavelet parameters for speed and accuracy. In particular, we used biorthogonal 3.1 wavelets with the Low Low High subband, a 15 × 15 pixel window with a step size of 5 pixels, and the co-occurrence matrix quantized into 15 gray levels with a 3 × 3 pixel displacement distance. The number of non-zero elements in the wavelet feature vector was then normalized to one to obtain the Wavelet OD. Droplets with a Wavelet OD between 0.3 and 0.7 were empirically identified as slow-growers and sorted into the keep channel. Droplets with a wavelet OD of less than 0.3 were typically empty droplets while droplets with a wavelet OD of greater than 0.7 contained a dense bacteria colony.

### Anaerobic chamber

Bacteria cultivation, droplet generation, and droplet sorting were all carried out inside a vinyl anaerobic chamber (Coy Laboratory Products) supplied with an 86% N_2_/10% O_2_/4% H_2_ gas mixture. The O_2_ and H_2_ concentrations were monitored using an anaerobic monitor (Coy CAM-12). The H_2_ concentration was maintained between 1.5-2.5% and the O_2_ concentration was typically less than 1 ppm. A hydrogen sulfide reducing column (Coy) was placed inside the chamber to prevent corrosion of the electronic components from H_2_S buildup.

### Media

#### BHIS

We first mixed 500 mL water, 18.5 g Brain Heart Infusion Broth (Sigma), 2.5 g yeast extract, 0.25 g L-cysteine, and 7.5 g agar, if making plates. We then autoclaved the solution and added 0.5 mL of 0.1% Vitamin K solution in 95% ethanol and 0.5 mL of a filter-sterilized hemin solution of 5 mg/mL in 0.1 M NaOH.

#### BHIS-ASF

For ASF cultures, we modified the BHIS media described above by adding 5% new born calf serum, 5% sheep serum, and 5% horse serum (all sera from Fisher Scientific).

#### GMM

GMM was prepared following the directions outlined by Goodman et al. (2011) [50].

#### YCFA

We used the following modified version of DSMZ media 1611. We first mixed 10 g casitone, 2.5 g yeast extract, 5 g dextrose (D-glucose), 0.045 g MgSO_4_ x 7 H_2_O, 0.09 g CaCl_2_ x 2 H_2_O, 0.45 g K_2_HPO_4_, 0.45 g KH_2_PO_4_, 0.9 g NaCl, 0.001 g resazurin sodium salt, 1.9 mL acetic acid, 0.7 mL propionic acid, 90 μL iso-butyric acid, 100 μL n-valeric acid, 100 μL iso-valeric acid, and 500 mL DI water. The media was then boiled for 10 min while stirring and then cooled. Next, we added 4 g NaHCO_3_, 0.5 g L-cysteine-HCl, 0.01 g hemin, 0.0025 g mucin, and 15 g agar, if making plates. DI water was then added to bring the total volume to 990 mL, the pH was adjusted to 6.7 – 6.8, and the solution was autoclaved. After autoclaving, we added 10 mL of filter-sterilized vitamin solution. One liter of vitamin solution is made by mixing 2 mg biotin, 2 mg folic acid, 12 mg pyridoxine-HCl, 4.5 mg thiamine-HCl, 5 mg riboflavin, 5 mg nicotinic acid, 5 mg D-Ca-pantothenate, 0.1 mg vitamin B12, 5 mg p-aminobenzoic acid, 5 mg DL-thioctic acid, and 1L DI water.

### ASF culture, DNA isolation, and amplification

ASF strains were grown in liquid broth and microfluidic droplets from both pure strain cultures and mixed community fecal cell suspensions. For pure strain cultures, each strain was grown in BHIS-ASF medium for 1 to 2 days and then droplets were generated in the microfluidic device by choosing an appropriate dilution factor. For mixed community cultures, stock fecal cell suspensions were first created by dissociating 3 ASF fecal pellets (Taconic Biosciences) in 5 mL of BHIS medium, letting the fecal solids sediment, collecting the supernatant, and storing at −80 °C in a 25% glycerol/25% BHIS/50% cell suspension solution. Droplets were generated from a 50x dilution of the mixed community stock solution. Both the liquid broth and droplet emulsions cultures were grown inside a 37 °C incubator. After cultivating ASF in droplets, we broke the emulsion by adding an equal part of 1H,1H,2H,2H-Perfluoro-1-octanol (PFO), vortexing and centrifuging the solution, and removing the oil and PFO. Finally, for both liquid broth and droplet cultures, we extracted the DNA using the Qiagen DNeasy Blood & Tissue Kit according to the manufacturer’s instructions with the exception that the total volume extracted from droplets was 100 μL. The 8 ASF strains were detected by PCR and droplet digital PCR (ddPCR) using species-specific primers previously developed [51]. We adapted the PCR protocol from Sarma-Rupavtarm et al. (2004) to detect the ASF strains present in the mixed community fecal samples before cultivation (see Supp. Fig. 2). We quantitatively determined the relative abundance of the 8 ASF species from the fecal cell suspension, liquid broth cultures, and droplet emulsion cultures using the BioRad QX200 Droplet Digital PCR System. For ddPCR, we mixed 10 µL QX200 ddPCR EvaGreen Supermix (BioRad), 200 nM forward and 200 nM reverse strain-specific primers (from 10 µM stock), and 9.2 µL of DNA-free water and extracted ASF DNA (where the extracted DNA dilution was first determined for each strain and sample). We then ran the ddPCR reaction mixtures through the QX200 Droplet Generator with Droplet Generation Oil for Evagreen (BioRad), thermocycled the ddPCR droplets for 10 min at 94 °C step followed by 35 cycles of 1 min at 94 °C, 1 min at 60 °C, and 2 min at 72 °C, and ran the thermocycled ddPCR droplets through the QX200 Droplet Reader. The copies/μL for each sample were determined by manually thresholding the fluorescence intensity reads. We then determined the relative abundance of each species within a sample by dividing the species’ counts by the total counts of all species within a sample.

### Human stool sample collection and droplet cultivation

In this study, we used a single stool sample collected previously from a fecal microbiota transplant donor [52]. Sample collection was approved by University of Chicago Ethics Committee and by the University of Chicago Institutional Review Board (IRB 132-0212) and written and informed consent was obtained for the stool donor. Stool aliquots were made by spinning down 50 μL of stool diluted in 100 μL of PBS, carrying forward the supernatant, and storing the supernatant at −80 °C. The live cell and dead cell densities in our aliquots were 1 × 10^9^/mL and 3 × 10^9^/mL, respectively (Live/Dead BacLight, ThermoFisher), indicating that bacteria viability was affected by the sample preservation process. For plate cultures, 100 μL of the aliquoted bacteria suspension was spread onto the agar plates. For droplets, aliquots were diluted 200x, so that according to Poisson statistics, ∼20-30% of droplets will contain one living bacteria. In a typical experiment, we generated and cultured a ∼0.5 – 1 mL emulsion of ∼65 – 115 pL droplets (i.e., ∼4 – 15 million droplets). After cultivation, the emulsion was broken and the DNA extracted following the same procedure as outlined above for ASF droplet cultures, with the following exception for the sorted droplets. Because our sorting rate was limited to ∼30 Hz, the total volume of the droplets which were sorted into the “keep” path was significantly less than 100 μL. In the two sorting experiments conducted here, the sorted droplets only formed a very thin layer on top of the oil. Roughly, the estimated total volume observed by pipetting, was 1 – 10 μL. To extract the DNA, we therefore first added 150 μL DNA-free water and 150 μL PFO to the sorted droplets followed by the same vortex and centrifugation steps.

In order to verify that the droplet cultivation platform is also compatible with traditional microbiology workflows, we further cultured bacteria grown in droplets on an agar plate after sorting. In one experiment, we cultured human stool bacteria in BHIS droplets for 1 day, sorted the droplets, and then streaked ∼10 μL of the sorted droplet emulsion and oil onto a BHIS agar plate. We then cultured the plate for 2 days at 37 °C and then randomly picked 24 colonies. Each colony was placed into 2 mL of BHIS broth, mixed, and 1 mL was extracted for Sanger sequencing. DNA was isolated using the Qiagen DNeasy Blood & Tissue Kit according to the manufacturer’s instructions. We amplified the 16S rRNA using the primers 27F (AGAGTTTGATCMTGGCTCAG) and 1492R (GGTTACCTTGTTACGACTT) by mixing 250 nM of the 27F and 1492R primers, 10 μL GoTaq Green Master Mix (Promega), 2 μL extracted DNA, and 7 μL DNA free water followed by running PCR amplification with (i) 3 min at 95 °C, (ii) 30 cycles of 30 sec at 95 °C, 30 sec at 54 °C, 1 min at 72 °C, and (iii) 10 min at 72 °C. Each sample was then run through a 2% agarose gel, the band was excised from the gel, and the DNA purified using the QIAquick Gel Extraction Kit (Qiagen) according to the manufacturer’s instructions. The DNA was then Sanger sequenced with the 27F and 1492R primers at the University of Chicago Comprehensive Cancer Center DNA Sequencing and Genotyping Facility. Finally, the forward and reverse reads were aligned in Benchling (https://benchling.com) using the MAFFT algorithm with default parameters and the consensus sequence was searched using the Standard Nucleotide BLAST for the closest taxonomic match.

### Sequencing

The Environmental Sample Preparation and Sequencing Facility at Argonne National Laboratory (Argonne, IL, USA) performed library preparation and sequencing of our DNA isolates. Briefly, 35 cycles of amplification were performed using the primer set described previously [53] that target the V4 region of the 16S rRNA gene to generate our amplicons from purified DNA, and Illumina MiSeq paired-end sequencing (2 × 300) was used to sequence our amplicon libraries. We analyzed the raw sequencing reads using illumine-utils [54] to (1) de-multiplexed raw sequencing reads into samples, (2) join paired-end sequences, and (3) remove low-quality sequences by requiring a minimum overlap size of 45 nucleotides between the two reads in each pair and removing any read that contained more than 2 mismatches in the overlapped region (mismatches in sequences survive these criteria were resolved with the use of the higher quality base). Finally, we inferred amplicon sequence variants (ASVs) in our dataset using Minimum Entropy Decomposition (MED) [32] through the oligotyping pipeline v2.1 [55], and the program Global Alignment for Sequence Taxonomy (GAST) assigned taxonomy to each ASV [56].

### Data Analysis

We characterized each sample using standard ecological metrics including richness (*R*), Shannon’s diversity index (*H’*), and rank-abundance curves. ASVs were first normalized by proportion so that for each sample, ∑*i p*_*i*_ = 1, where *p*_*i*_ is the proportional abundance of ASV *i*. The richness is the total count of ASVs detected within a sample and Shannon’s index, *H’* = -∑*i p*_*i*_ log (*p*_*i*_). Rank-abundance curves were obtained by ordering the ASVs by decreasing *p*_*i*_ for each sample. Additionally, we also counted the richness of ASVs for each sample which were <1% abundant in the raw stool sample (*R*_*low*_) in order to investigate if droplets can enhance the cultivation of rare species. We statistically tested the metrics *R, H’*, and *R*_*low*_ using the nonparametric Mann-Whitney *U* test under the null hypothesis that the difference between the means of the metrics on plates and droplets, irrespective of culture media and cultivation time, is zero. The statistical testing was implemented in the R package using the function wilcox.test. We also statistically tested if there was any difference in the rank-abundance distributions between samples with the same cultivation condition (i.e., droplet or plate) and the same media. In particular, we applied the Kolmogorov-Smirnov test (R function ks.test) under the null hypothesis that two samples can be generated from the same distribution. Next, hierarchical clustering was applied to infer associations between samples. Samples were clustered at the family level by calculating the Bray-Curtis dissimilarity index in the R function vegdist in the package vegan and then plotting the dendrogram. Finally, we statistically tested that *Bacteroides* oligotypes cluster closer to raw stool using hierarchical cluster analysis with multiscale bootstrap resampling (R function pvclust) [57].

We also tested if sorting slow-growing colonies could amplify the relative abundance of rare ASVs. Rank-abundance curves for the two independent sorting experiments were first generated and the combined sorted and unsorted distributions were statistically compared using the two-sample Kolmogorov-Smirnov test, which tests if two sample distributions can be drawn from a common distribution. Next, we investigated the ability of the sorted droplets to increase the abundance of low-abundant ASVs from the raw stool. We arbitrarily set the limit to 1%; ASVs which were <1% abundant in raw stool were considered amplified if the ASV’s relative abundance was >1% in the sorted or unsorted droplets. The ASVs which satisfied this condition across two separate sorting experiments in BHIS droplets were pooled together for phylogenetic analysis. The phylogenetic tree was generated by performing multiple sequence alignment using the default settings on Clustal Omega followed by calculating a DNA Neighbour Joining tree in Jalview. Finally, the increase in proportional abundance for the ASVs which satisfied the above condition in unsorted and sorted droplets was calculated.

We used anvi’o v5.5 [58] to visualize heat map visualizations of ASV percent relative abundances and clustering dendrograms, and used the open-source vector graphics editor Inkscape (available from https://inkscape.org) to finalize them for publication.

## Acknowledgements

We thank Dr. Michael Wannemuehler for generously providing the 8 pure ASF strains and for notes on culturing the ASF, the Environmental Sample Preparation and Sequencing Facility at Argonne National Laboratory, the Pritzker Nanofabrication Facility, and the University of Chicago Comprehensive Cancer Center DNA Sequencing and Genotyping Facility. This research is supported by the NIDDK P30 University of Chicago Digestive Disease Research Core Center (DK42086), T32 DK07074 (supporting WJW), RC2 DK122394-01, The Samuel and Emma Winters Foundation Grant (2018-2019), and the GI Research Foundation of Chicago (supporting WJW). We also thank James & Katie Mutchnik (who supported AME) and the Duchossois Family Institute at the University of Chicago (for pilot funding to WJW, MT, AME, and ST).

