## Supplementary material for "High-throughput isolation and sorting of gut microbes reduce biases of traditional cultivation strategies": video 1

### Supplementary Information

#### Supplementary Table File Descriptions

1. **Experiment_Info.** The droplet generation and cultivation information is listed for each experiment.
2. **ASF_Copies_Per_uL.** Manually thresholded ddPCR copies/µL for each ASF species and experiment.
3. **ASV_Counts_Matrix.** The counts matrix of each ASV determined by Minimum Entropy Decomposition (MED) for the raw human stool sample and the human stool cultured on plates and in droplets.
4. **Sequences_Taxonomy.** Closest taxonomic assignment of each ASV from raw and cultivated human stool samples.
5. **Sequences.** Sequences from MED analysis for raw and cultivated human stool samples.
6. **Oligotyping_Percent_Matrix.** Percent matrix for *Bacteroides* oligotype assignment for each sample.
7. **Oligotyping_Representatives.** Oligotypes of *Bacteroides spp.*
8. **Sanger_Consensus_Sequences.** The consensus sequence for each of 24 randomly picked colonies from sorted droplets which were streaked onto an agar plate. For each sequence, the closest BLAST taxonomic assignment along with percent identity is also listed.

#### Supplementary Video Descriptions

1. **Video_1.** Human stool bacteria cultured in BHIS droplets for 1 day.
2. **Video_2.** Droplet sorting in an anaerobic environment for human stool bacteria cultured in BHIS droplets for 1 day. The top left time stamp is in seconds. For each droplet, the region between the 2 red dots is analyzed using the Wavelet OD. For empty droplets and droplets containing a dense colony, the Wavelet OD does not satisfy the thresholding criteria (the decision is labeled as false, ‘F’) and the droplets flow down the waste path. When the threshold criteria is met for a sparse colony (the decision is labeled as true, ‘T’), the electrodes are actuated sending the droplet to the ‘keep’ path. We note that the spots in the oil phase were observed even for droplets generated without bacteria.

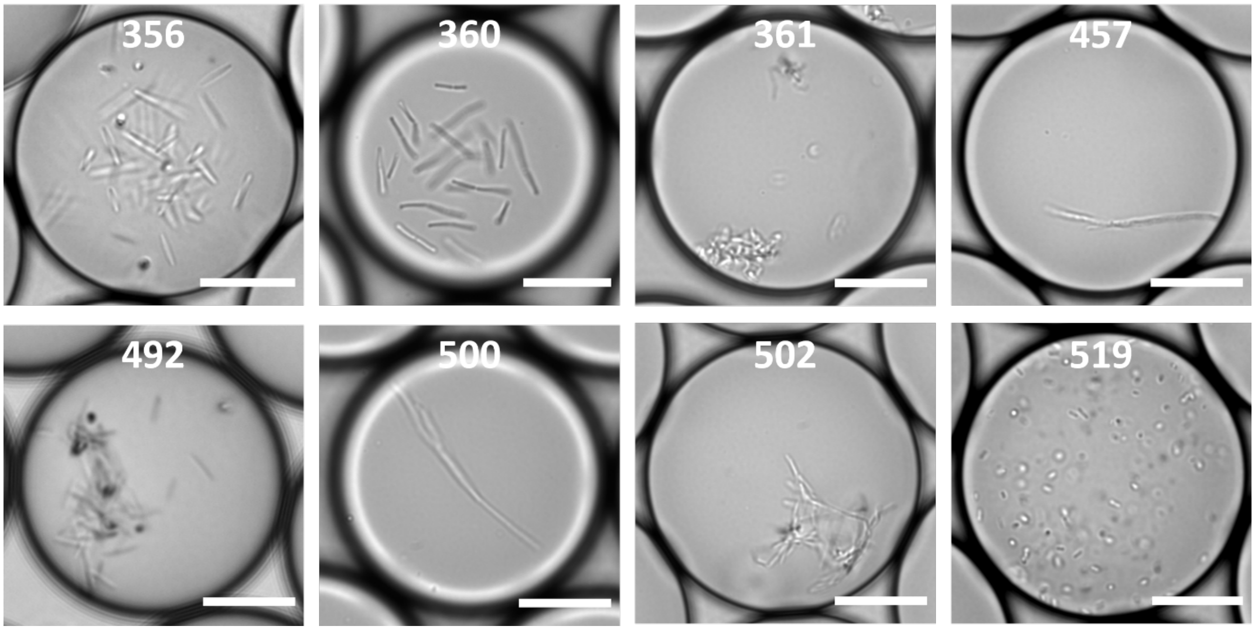


**Supplementary Figure 1.** Typical colony morphologies of the 8 ASF strains grown in microfluidic droplets after 17 to 20 hours. Colony morphologies differ drastically between species in the same genus, i.e., *Lactobacillus intestinalis* (ASF 360) and *Lactobacillus murinus* (ASF 361), and also *Clostridium* sp. (ASF 356) and *Clostridium* sp. (ASF 502). Scale bars are 20 µm.

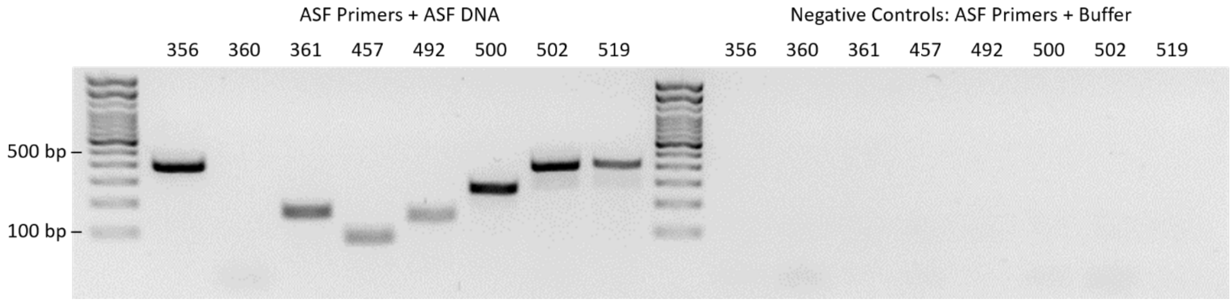


**Supplementary Figure 2.** All ASF species were detected in a fecal pellet by PCR using species specific primers except ASF 360. The PCR protocol is as follows. ASF DNA was extracted from the fecal pellet cell suspension using the Qiagen DNeasy Blood and Tissue kit following the manufacturer's instructions. The measured DNA concentration using NanoDrop was 10.4 ng/µL. The following PCR reagents were first mixed: 10 µL GoTaq Green Master Mix (Promega), 4.2 µL nuclease free water, 200 nM forward and 200 nM reverse strain-specific primers (Sarma-Rupavtarm et al., 2004), and 5 µL of ASF DNA (left column set) or 5 µL AE buffer (right column set, negative controls). PCR amplification was then performed with a 10 min, 94 °C step followed by 35 cycles of 1 min at 94 °C, 1 min at 60 °C, and 2 min at 72 °C. The PCR products were mixed with a gel loading dye and ran in a 2% agarose gel with 1x SYBR Safe DNA Gel Stain. The expected length for each species' product is: ASF 356 (413 bp), ASF 360 (132 bp), ASF 361 (182 bp), ASF 457 (95 bp), ASF 492 (167 bp), ASF 500 (284 bp), ASF 502 (427 bp) and ASF 519 (429 bp). No bands are seen in the negative controls.

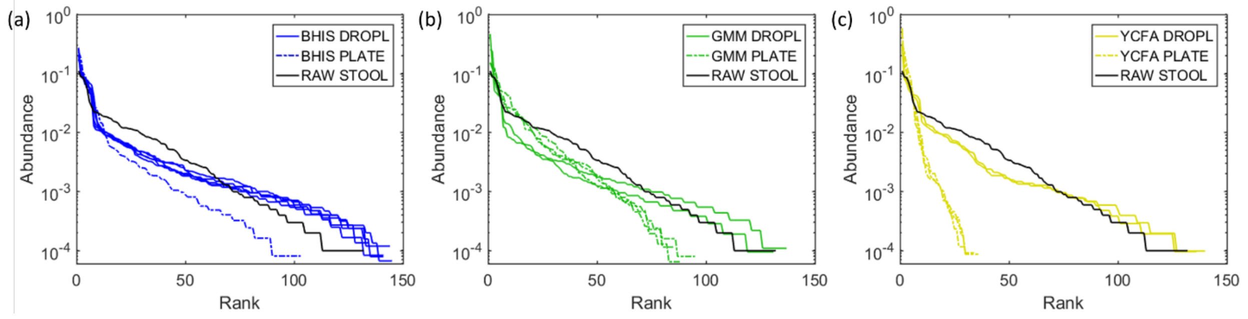


**Supplementary Figure 3.** Rank-abundance curves for independent experiments in BHIS, GMM, and YCFA media cultivated in droplets or on agar plates. No two experiments within a given media and condition (i.e., BHIS droplet, GMM droplet, GMM plate, YCFA droplet, or YCFA plate) were found to reject the null hypothesis that the two data samples can be generated from the same distribution under the two-sample Kolmogorov–Smirnov test, indicating the reproducibility of cultivation between independent experiments.

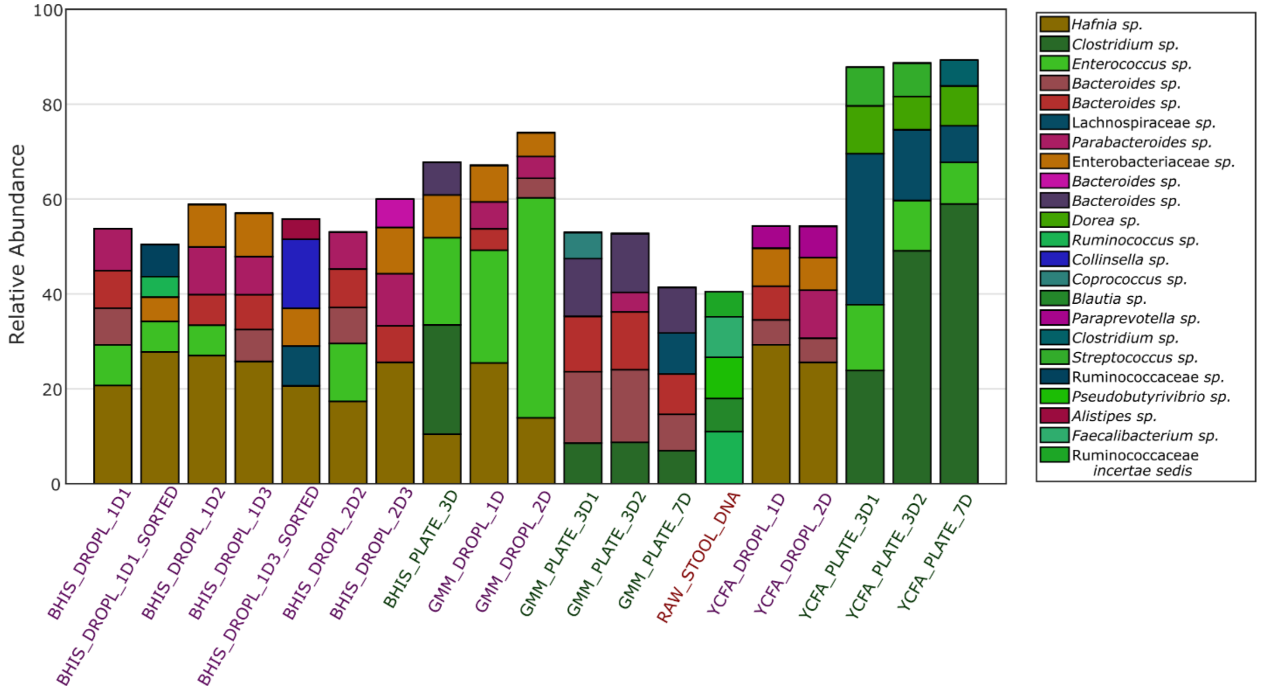


**Supplementary Figure 4.** Relative abundance of the top 5 most abundant ASVs in each sample with taxonomic identification.

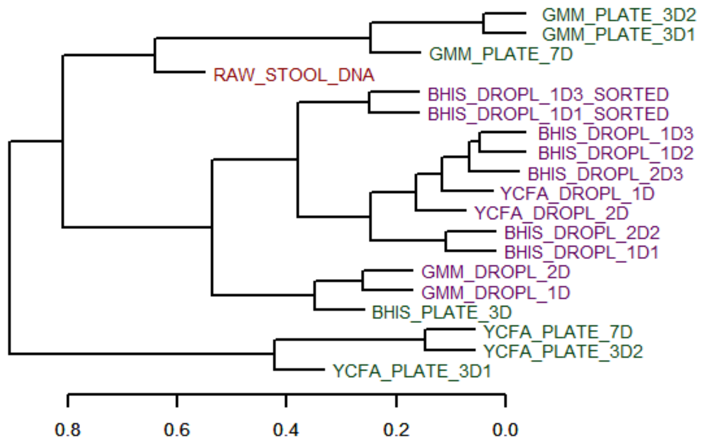


**Supplementary Figure 5.** Bray-Curtis hierarchical clustering of the family-level composition for all samples. The droplet-based culture reduces the variation in cultivated composition between media as compared to agar plate cultures.

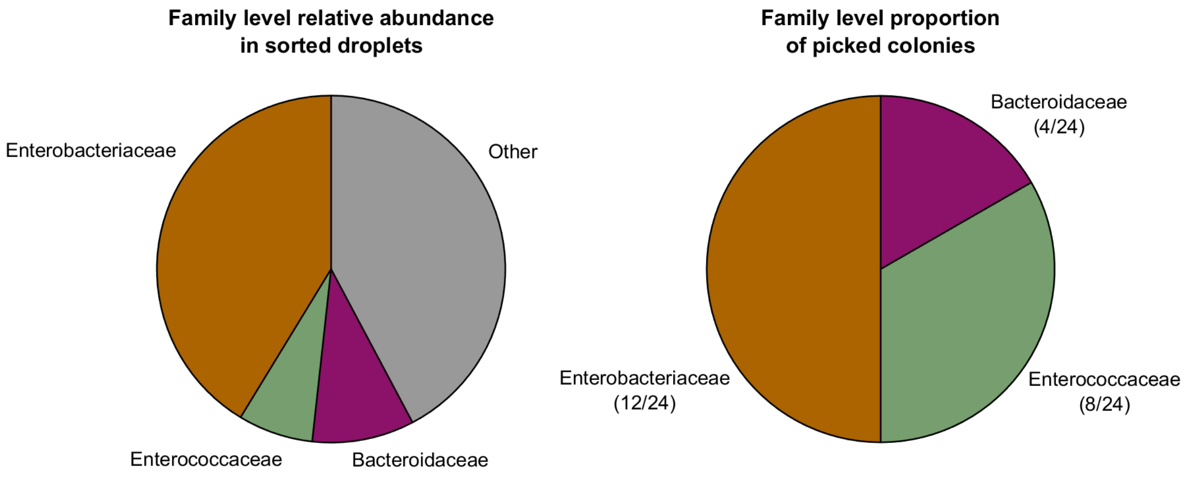


**Supplementary Figure 6.** Bacteria can be cultured on traditional agar plates following cultivation in droplets. (Left) The family level relative abundance of the pooled droplets from experiment BHIS_DROPL_1D1_SORTED. (Right) The family level taxonomic assignment of 24 randomly picked colonies on a BHIS agar plate streaked from the same sample shown on the left. For each family detected on agar plate, only one genera was observed within that family. In particular, we detected *Hafnia* (12/24), *Enterococcus* (8/24), and *Bacteroides* (4/24).
